# Multi-session visual cortex transcranial random noise stimulation in adults with amblyopia

**DOI:** 10.1101/679662

**Authors:** Richard Donkor, Andrew E. Silva, Caroline Teske, Margaret Wallis-Duffy, Aaron Johnson, Benjamin Thompson

## Abstract

**Purpose:** We tested the hypothesis that five daily sessions of visual cortex transcranial random noise stimulation would improve contrast sensitivity, crowded and uncrowded visual acuity in adults with amblyopia.

**Methods:** Nineteen adults with amblyopia (44.2 ± 14.9yrs, 10 female) were randomly allocated to active or sham tRNS of the visual cortex (active, n = 9; sham, n = 10). tRNS was delivered for 25 minutes across five consecutive days. Monocular contrast sensitivity, uncrowded and crowded visual acuity were measured before, during, 5 minutes and 30 minutes post stimulation on each day.

**Results:** Active tRNS significantly improved contrast sensitivity and uncrowded visual acuity for both amblyopic and fellow eyes whereas sham stimulation had no effect. An analysis of the day by day effects revealed large within session improvements on day 1 for the active group that waned across subsequent days. No long-lasting (multi-day) improvements were observed for contrast sensitivity, however a long-lasting improvement in amblyopic eye uncrowded visual acuity was observed for the active group. This improvement remained at 28 day follow up. However, between-group differences in baseline uncrowded visual acuity complicate the interpretation of this effect. No effect of tRNS was observed for amblyopic eye crowded visual acuity.

**Conclusions:** In agreement with previous non-invasive brain stimulation studies using different techniques, tRNS induced short-term contrast sensitivity improvements in adult amblyopic eyes, however, multiple sessions of tRNS did not lead to enhanced or long-lasting effects for the majority of outcome measures.

## Introduction

Amblyopia is a developmental disorder of the visual cortex, with a prevalence of approximately 1-5% [1–3]. Amblyopia causes a wide range of vision deficits [4,5] including a monocular loss of high-contrast visual acuity [6,7] that is particularly pronounced for crowded optotypes [8,9], reduced contrast sensitivity in the affected eye [6,10–12], and impaired or absent stereopsis [13,14]. Amblyopia is also associated with chronic suppression of the affected eye [15,16] that may play a key role in the etiology of the disorder [17].

Amblyopia involves abnormal processing within the primary and extrastriate visual cortex [18] and therefore recovery from amblyopia requires a change in cortical function. Current amblyopia treatments achieve this by directly manipulating visual input to the brain. For example, the most common amblyopia treatment involves the provision of a clear retinal image in the amblyopic eye using refractive correction followed by occlusion of the non-amblyopic eye. This treatment improves amblyopic eye visual acuity, but has drawbacks in terms of compliance [19] and reduced efficacy with increasing age [20].

Transcranial electrical stimulation (tES) refers to a suite of non-invasive neuro-modulation techniques including transcranial direct current stimulation (tDCS), transcranial alternating current stimulation (tACS) and transcranial random noise stimulation (tRNS) that may enhance plasticity in targeted regions of the human brain [21–23], including the visual cortex [24–26]. Currently, tES methods are being investigated as a potential neurorehabilitation tool for disorders including stroke [27–30], chronic pain [31,32] and tinnitus [33,34] and there is growing interest in the use of tES to treat disorders of vision (see [35,36] for recent reviews).

Following early work that reported improved contrast sensitivity in adults with amblyopia after visual cortex transcranial magnetic stimulation [37,38], a number of studies have investigated the application of anodal tDCS to amblyopia. A single session of anodal tDCS improves amblyopic eye contrast sensitivity [39,40], increases visually evoked potential (VEP) amplitudes induced by stimuli presented to the amblyopic eye [40], and balances the response to inputs from each eye within visual cortex [39]. Furthermore, anodal tDCS enhances the effect of perceptual learning (PL) in adults with amblyopia [26] and recent studies have revealed that visual acuity, detection thresholds, and stereopsis improve in mature amblyopic rats following anodal tDCS [41–43]. One potential mechanism for anodal tDCS effects in adults with amblyopia is a reduction in GABA-mediated inhibition [44] within the visual cortex. GABA has been associated with interocular suppression in strabismic cats [45] and may act as a “break” on visual cortex plasticity [46].

A recently developed tES technique, tRNS, involves an alternating current that randomly changes in frequency and amplitude [47]. tRNS may have larger effects on cortical activity than other tES protocols. For example, tRNS induced significantly greater improvements in tinnitus symptoms [33] and larger increases in motor evoked potential amplitude (MEP) [21] compared to either tDCS or tACS. Furthermore, visual cortex tRNS enhanced visual perceptual learning in adults with normal vision to a greater extent than anodal tDCS [48,49] and the combination of tRNS and perceptual learning enhanced the transfer of learning to non-trained visual tasks in adults with amblyopia [24]. tRNS has also been reported to enhance visual perceptual learning in adults with cortical blindness [49]. Potential mechanisms for these effects include an acute enhancement of the signal-to-noise ratio within the visual cortex due to stochastic resonance [50–52] and longer-lasting alterations in neural membrane function due to repetitive opening and closing of sodium channels [53].

Within this single-blind, between subjects, randomized, sham-controlled study, we tested the hypothesis that five daily sessions of visual cortex tRNS alone would lead to improved amblyopic eye contrast sensitivity, crowded and uncrowded amblyopic eye visual acuity in adult patients. We chose not to combine tRNS with visual perceptual learning because the effects of multi-session visual cortex tES alone have not yet been characterized in adults with amblyopia. Furthermore, visual perceptual learning is intensive and can be challenging to deploy in a clinical environment. We reasoned that if multi-session tRNS alone could improve a range of visual functions in adults with amblyopia, it may represent a potential treatment approach in its own right.

## Methods and materials

### Participants

Nineteen healthy adults with amblyopia (mean age 44.2 ± 14.9 yrs) participated (**Table 1**). Participants were randomly assigned to the active tRNS group or to the control (sham) group. Amblyopia was defined as reduced best corrected visual acuity (> +0.3 logMAR) in one eye in the absence of ocular pathology and at least a 0.2 logMAR acuity difference between the eyes. Anisometropia was defined as a difference in spherical equivalent between the two eyes of ≥ 0.50 Diopters (D), or a difference of astigmatism in any meridian ≥ 1.50 D [54]. Initial visual acuity was measured using an M&S Snellen chart. Inclusion criteria were: (i) presence of strabismic and/or anisometropic amblyopia; (ii) 0.0 logMAR visual acuity or better in the fellow eye (FE). Exclusion criteria [55–58] were: (i) presence of a scalp skin condition that contraindicated tRNS; (ii) history of neurological or psychiatric disorders, such as seizures; (iii) current medication for the treatment of neurological or psychiatric disorders; (iv) a history of brain injury; (v) implanted medical devices. Data were collected within a clinical optometric practice to provide a real-world clinical setting. Potential participants were contacted following a search of the clinic’s patient database. Interested participants completed telephone screening to determine eligibility. The experimental procedures were approved by the Ethics Review Board of the University of Waterloo, Canada and were consistent with the declaration of Helsinki. Written informed consent was obtained from all participants. All participants were remunerated for their time. Two subjects were excluded due to loss of follow up after the first day (**Figure 1**).

**Table 1:** Clinical details of the study participants **(AG:** Active group; **SG:** Sham group; **AME:** Amblyopic eye; **FFE:** Fellow fixating eye; **ADD:** Near power addition; **Anise:** Anisometropia; **Strab:** Strabismus; **Mixed:** Anisometropia & Strabismus; **M:** Male; **F:** Female).

| Identification | Age/sex | Patching | Type of amblyopia | Stereoaucuity | Visual acuity (logMAR) |  | Current refraction |  | ADD |
| --- | --- | --- | --- | --- | --- | --- | --- | --- | --- |
|  |  |  |  |  | AME | FFE | AME | FFE |  |
| AG1 | 27/M | Yes | Mixed | <800 | +1.40 | 0.00 | +2.0 -0.50 x 155 | +0.50 DS |  |
| AG2 | 20/M | Yes | Aniso | 400 | +0.50 | -0.05 | +1.00 -1.50 x 175 | Plano |  |
| AG3 | 37/M | Yes | Aniso | <800 | +0.30 | 0.00 | -1.00 -1.00 x 100 | -0.25 -2.50 x 090 |  |
| AG4 | 45/F | Yes | Aniso | 200 | +0.30 | 0.00 | +2.75 -0.75 x 080 | +2.00 -0.25 x 159 | +1.50 |
| AG5 | 59/M | No | Aniso | <800 | +0.70 | 0.00 | +8.25 DS | +8.25 -1.25 x 105 | +2.50 |
| AG6 | 41/F | Yes | Mixed | <800 | +0.50 | 0.00 | +2.00 DS | +0.25 -3.75 x 179 | +0.75 |
| AG7 | 46/M | No | Aniso | <800 | +0.40 | 0.00 | +0.75 DS | +4.25 -0.50 x 165 |  |
| AG8 | 52/F | No | Aniso | 400 | +0.70 | 0.00 | +0.50 -0.50 x 175 | +2.00 -0.50 x 075 | +2.25 |
| AG9 | 21/F | No | Aniso | 60 | +0.50 | +0.10 | +0.50 -4.00 x 180 | Plano |  |
| SG1 | 51/F | Yes | Mixed | <800 | +0.30 | -0.09 | +3.25 -0.50 x 165 | Plano | +2.25 |
| SG2 | 21/F | No | Aniso | <800 | +0.30 | 0.00 | +4.50 -2.50 x 180 | +3.50 - 1.50 x 167 |  |
| SG3 | 58/F | No | Aniso | <800 | +0.50 | 0.00 | +4.50 -3.75 x 176 | +3.75 -3.75 x 176 | +2.25 |
| SG4 | 58/M | Yes | Aniso | 400 | +0.30 | 0.00 | +4.00 -0.25 x 180 | +1.50 -0.50 x 180 | +2.75 |
| SG5 | 52/F | No | Aniso | 200 | +0.40 | 0.00 | -2.50 -0.50 x 155 | -1.50 DS |  |
| SG6 | 19/M | No | Aniso | <800 | +0.30 | 0.00 | +2.25 DS | +0.75 DS |  |
| SG7 | 55/M | No | Strab | <800 | +0.30 | 0.00 | +1.75 -0.75 x 085 | +1.75 -0.75 x 090 | +2.00 |
| SG8 | 49/M | No | Strab | 100 | +0.30 | 0.00 | +1.75 DS | +1.50 DS |  |
| SG9 | 24/M | No | Mixed | <800 | +0.30 | 0.00 | +8.25 -2.00 x 160 | +6.50 -1.25 x 0.40 |  |
| SG10 | 58/F | No | Strab | <800 | +0.50 | 0.00 | +4.50 -0.25 x 176 | +3.50 -0.50 x 165 | +2.75 |

**Figure 1:**
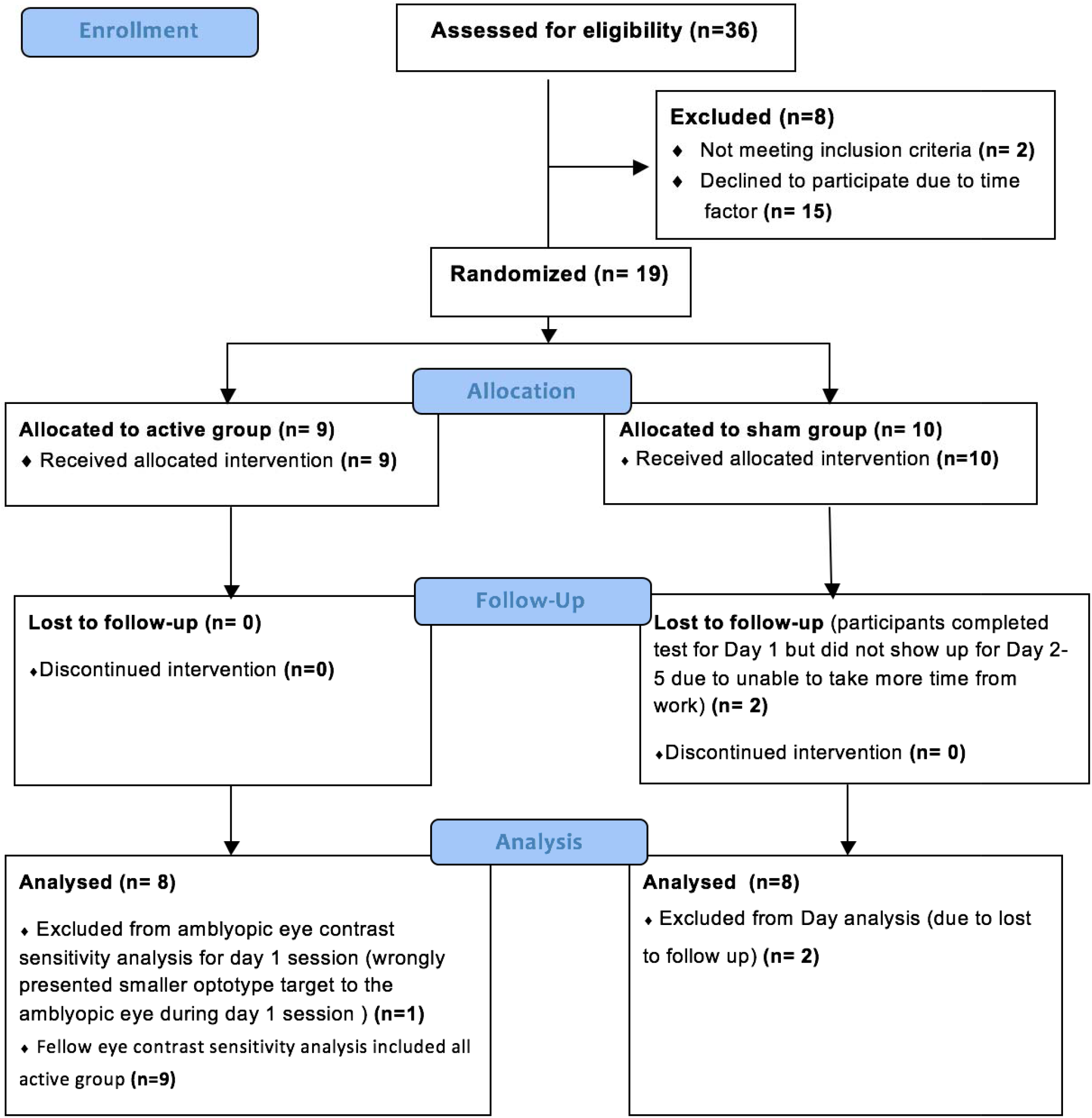
CONSORT flow diagram for the study

## Procedure

A single-blind, sham controlled, between-subjects design was adopted. Randomization followed allocation concealment procedures and was conducted by an experimenter who was not involved in data collection or eligibility assessment using a random number generator. Randomization occurred after participants had met the eligibility criteria and completed study enrolment. Participants completed 5 consecutive daily stimulation sessions and a follow-up session 28 days after the final stimulation session. Outcome measures were completed by the participants using automated computer programs with no input from the experimenter to minimize experimenter bias as much as possible.

### Transcranial Random Noise Stimulation

Subjects either received tRNS (2.0 mA, current density: 0.08 mA/cm, frequency range 0.1-640 Hz) or placebo (sham) stimulation of the primary visual cortex for 25 min over 5 consecutive days. Stimulation was delivered using a DC-stimulation MC device (Eldith, NeuroConn GmbH, Germany). There is no statistically significant difference between the effect of high frequency (101-640 Hz) and low frequency (0.1-100 Hz) visual cortex tRNS on visual perceptual learning enhancement [48]. Therefore, we chose to deliver the full frequency range. The stimulation was delivered via a pair of saline-soaked surface sponge electrodes (5 cm × 5 cm, 25 cm^2^ placed at Cz and Oz [59], as determined by the international 10/20 electroencephalogram system. The AC current was initially ramped up to a maximum of 2mA over 30 s and ramped down to 0mA over 30 s at the end of the stimulation session. During sham stimulation, the 30 s ramp-up was immediately followed by the ramp-down out [60]. Our between subjects design and use of participants entirely naïve to non-invasive brain stimulation ensured that participants remained masked to their treatment allocation. Our application of tRNS conformed to tDCS safety guidelines [58,61].

After the final tRNS sessions, participants were asked to rate the following sensations on a four-level scale (none, mild, moderate and severe): headache, neck pain, scalp pain, tingling, itching, burning sensation, sleepiness, trouble concentrating and acute mood change. Participants were also asked to rate whether any reported sensations were due to tES by selecting from the following options: no, remote chance, probable, definitely.

### Visual Function Measurements

Monocular contrast sensitivity and visual acuity (both crowded and uncrowded) were measured for each eye before, during, 5 min post, and 30 min post stimulation on each stimulation day (**Figure 2**). All measurements were made using Landolt-C optotypes presented using the Freiburg Vision Test (‘FrACT’) [62,63] software package on a MacBook Pro (Version 10.13.6, 13-inch, 2.7 GHz, 2560 × 1600). Gamma correction was conducted using a Spyder photometer and the FrACT software provided 10 bits of contrast resolution The Landolt-C optotype was presented at 8 possible orientations and viewed from 3 m in a dark room. Participants identified the gap orientation using button presses. Trials were self-paced with a maximum display time of 30 s. A Bayesian adaptive (“Best PEST”) algorithm controlled optotype size for the crowded and uncrowded visual acuity threshold measurements and optotype contrast for the contrast sensitivity threshold measurement. Each threshold measurement lasted approximately 3 mins. For the during stimulation condition, threshold measurement started 5 minutes into the stimulation. Landolt-C gap width was fixed at 30 arcmin for the non-amblyopic eye and 100 arcmin for the amblyopic eye during the contrast sensitivity measures. These parameters were based on pilot observations in individuals with moderate and severe amblyopia who could not resolve the 30 arcmin stimuli. Crowded optotypes were surrounded by a circle. Both the fellow eye and amblyopic eye were tested monocularly with the fellow eye tested first within each block. Uncrowded visual acuity was tested first within each block followed by crowded visual acuity and contrast sensitivity.

**Figure 2:**
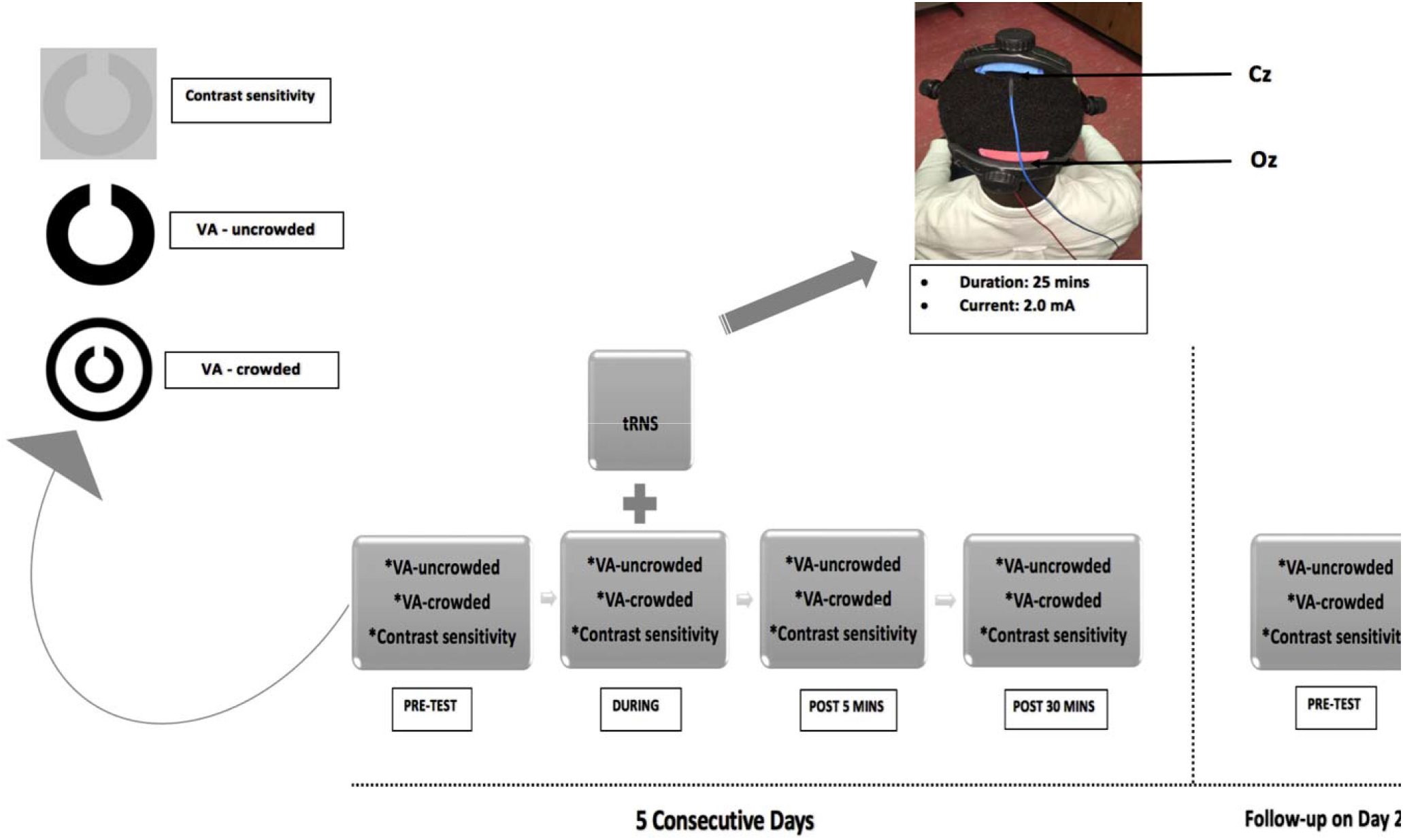
Testing and stimulation protocol. Each measurement was recorded before stimulation (pre-test), during stimulation, 5 min after stimulation (Post 5 mins) and 30 min after stimulation (Post 30 mins) for 5 consecutive days (middle column). Baseline (pre-test) measurements were recorded again for each eye 28 days (Day 28) after the last stimulation session. Stimulation was delivered for 25 mins at 2.0mA (right column). Active and reference electrodes were placed at Oz and Cz respectively. VA = visual acuity.

### Data Analysis

Statistical analyses were performed in R (R Core Team, 2020) [64] using the Bootstrap-Coupled Estimation package [65]. Visual acuities were recorded in logMAR units. Contrast sensitivity was recorded in log units. To test for tRNS effects within the 5 stimulation sessions, a mixed-effects analysis of variance (ANOVA) with a between-subjects factor of Group (active vs sham), a within-subjects factor of Day (day 1-5), and a within-subjects factor of Time (baseline, during, post 5 and post 30 mins) was conducted for each measurement type for each eye separately. Planned pairwise comparisons (least significance difference test) between baseline and all other timepoints were examined for each day. In addition, to assess whether tRNS had cumulative or long-term effects on visual function, a mixed ANOVA with factors of Group (active vs sham) and Baseline (baseline day 1, baseline day 2, baseline day 3, baseline day 4, baseline day 5, baseline day 28) was conducted for each outcome measure for each eye. All ANOVA ranalyses eported passed Levene’s test for equality of variances (p>.05) and test of sphericity (p>.05). Pairwise comparisons were conducted using the effect size Hedge’s *g* by bootstrap estimation (5000 bootstrap samples with replacement), with the 95% confidence interval around the *g* being bias-corrected and accelerated [65]. The permutation *P* values reported are calculated with 5000 reshuffles of the baseline and test labels performed for each permutation, with the P value indicating the likelihood of observing the mean difference, if the null hypothesis of zero difference is true, at an α of .05. Between group differences in the strength of any sensations induced by tRNS were assessed using the chi-squared test.

## Results

There were no adverse effects of tRNS, and there were no statistically significant between-group differences in the range or severity of subjective sensations reported (**Table 2**). Two participants in the sham group withdrew from the study after day 1 due to the time commitment required by the study and were excluded from the analysis (**Figure 1**). A technical error prevented an accurate amblyopic eye contrast sensitivity measurement on day 1 for one participant in the active group. This participant was excluded from the amblyopic eye contrast sensitivity analysis only (**Figure 1**).

**Table 2:** Subjective experiences reported by participants after the day 5 active or sham tRNS session.

| | None | Mild | Moderate | Severe | Chi-Square ($X^2$) |
| --- | --- | --- | --- | --- | --- |
| **Headache** |  |  |  |  | 0.060 |
| Active (N=9) | 4 | 1 | 4 | 0 |  |
| Sham (N=10) | 8 | 2 | 0 | 0 |  |
| **Neck pain** |  |  |  |  | 0.466 |
| Active (N=9) | 7 | 2 | 0 | 0 |  |
| Sham (N=10) | 9 | 1 | 0 | 0 |  |
| **Scalp pain** |  |  |  |  | 0.279 |
| Active (N=9) | 8 | 1 | 0 | 0 |  |
| Sham (N=10) | 10 | 0 | 0 | 0 |  |
| **Tingling** |  |  |  |  | 0.460 |
| Active (N=9) | 2 | 4 | 3 | 0 |  |
| Sham (N=10) | 3 | 6 | 1 | 0 |  |
| **Itching** |  |  |  |  | 0.277 |
| Active (N=9) | 3 | 4 | 2 | 0 |  |
| Sham (N=10) | 5 | 5 | 0 | 0 |  |
| **Burning sensation** |  |  |  |  | 0.074 |
| Active (N=9) | 4 | 2 | 3 | 0 |  |
| Sham (N=10) | 9 | 1 | 0 | 0 |  |
| **Sleepiness** |  |  |  |  | 0.751 |
| Active (N=9) | 6 | 2 | 1 | 0 |  |
| Sham (N=10) | 5 | 3 | 2 | 0 |  |
| **Trouble concentrating** |  |  |  |  | 0.330 |
| Active (N=9) | 9 | 0 | 0 | 0 |  |
| Sham (N=10) | 9 | 1 | 0 | 0 |  |
| **Acute mood change** |  |  |  |  | 0.330 |
| Active (N=9) | 9 | 0 | 0 | 0 |  |
| Sham (N=10) | 9 | 1 | 0 | 0 |  |
| **Others** | None | Jaw stiffness | Neck tension |  | 0.289 |
| Active (N=9) | 7 | 1 | 1 |  |  |
| Sham (N=10) | 10 | 0 | 0 |  |  |
| **Any of the symptoms related to tES** | None | Remote | Probable | Definite | 0.073 |
| Active (N=9) | 0 | 0 | 3 | 6 |  |
| Sham (N=10) | 2 | 2 | 0 | 6 |  |

### Contrast Sensitivity

For the amblyopic eyes (**Figure 3** - upper panel), there was a significant interaction between Group and Time, F_3,42_ = 3.584, p = .022, η_p_^2^ = .216. No other omnibus main effects or interactions were significant. Planned pairwise comparisons between baseline and all other timepoints were examined for the active and sham groups for each day. During day 1, the active group exhibited a significant improvement in contrast sensitivity from baseline for all post-test measurements (during: *g* =.272 [.195, .597], p = 0.01; post 5 min: *g* =.236 [.039, .726], p = .035; and post 30 min: *g* = .438 [.052, 1.207], p = 0.034: **Figure 4)**. No significant differences between baseline and any post-test were found for days 2-5. No significant differences between baseline and any post-test were found within the sham group for any day.

**Figure 3:**
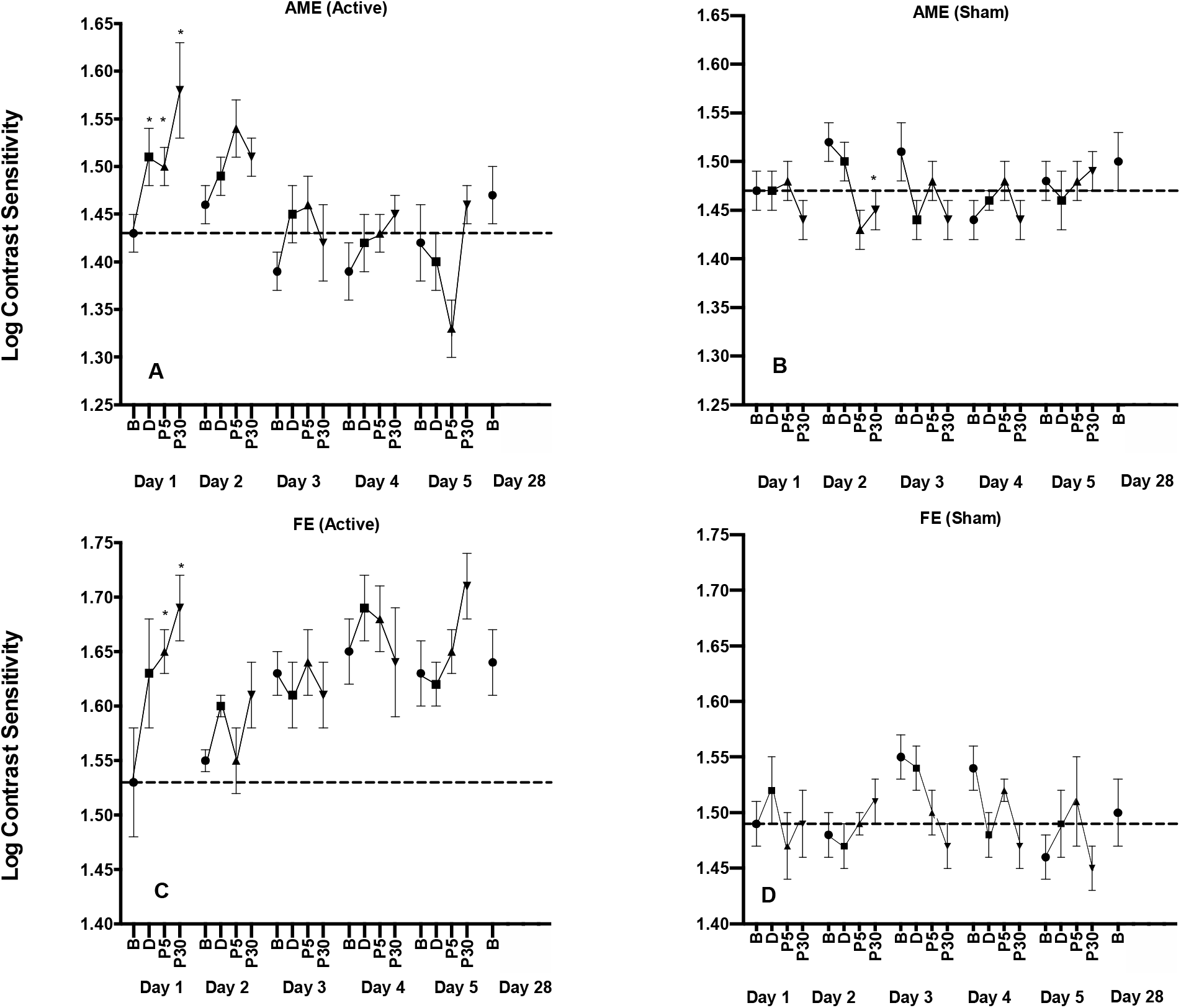
The effects of tRNS on contrast sensitivity during each daily session and at the day 28 follow-up visit. Data are shown separately for the amblyopic (top row) and fellow (bottom row) eyes and for the active (left column) and control (right column) groups at baseline (B) and during (D), 5 min (P5) and 30 min (P30) post tRNS. *Statistically significant difference from baseline (p < 0.05). Error bars show within-subject standard error of the mean (SEM). The dashed horizontal lines represent the mean before-stimulation threshold on day 1. Larger y-axis values indicate better contrast sensitivity.

**Figure 4:**
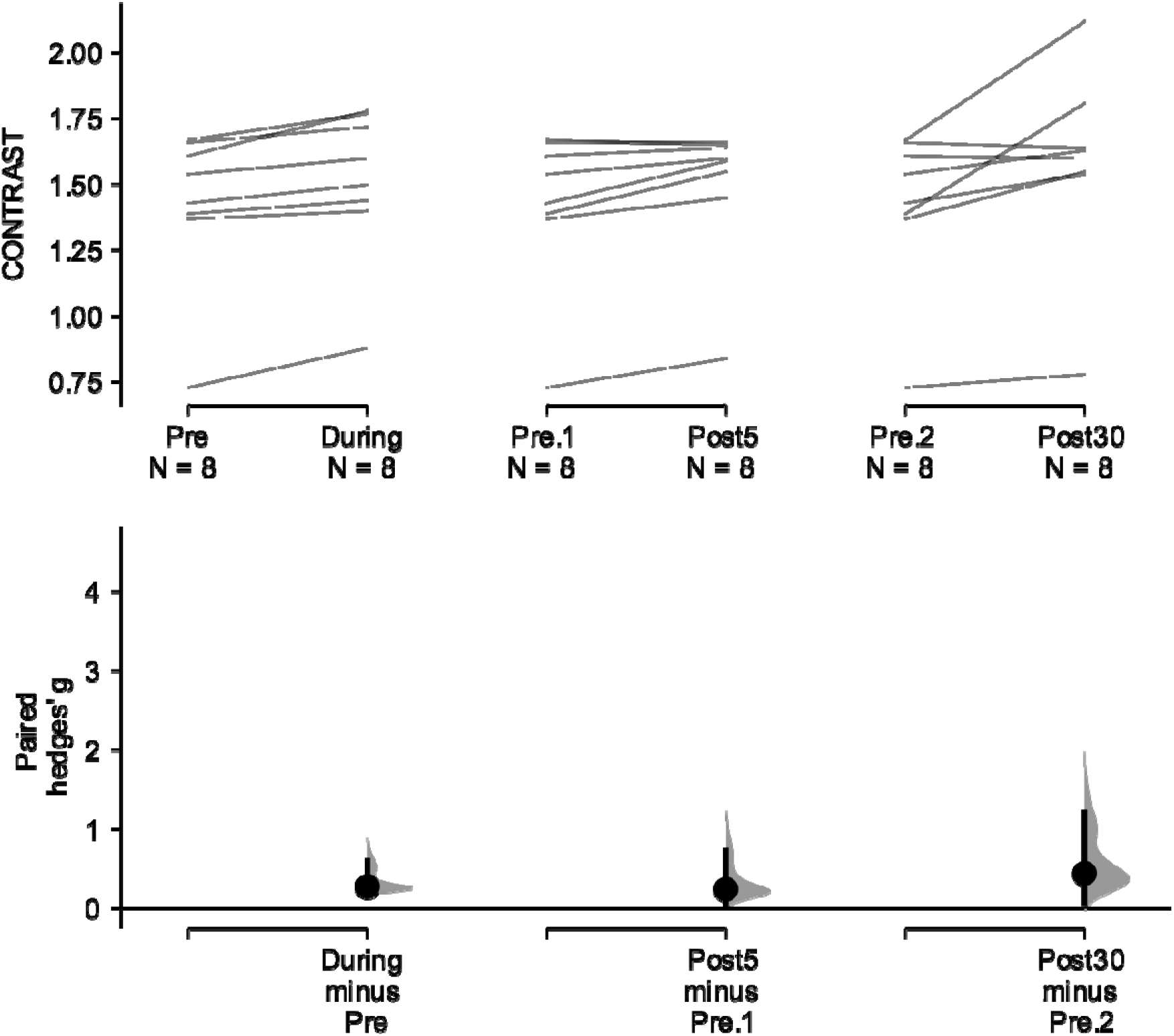
Paired Hedges’ *g* for three comparisons (during stimulation, Post 5 mins, Post 30 mins) to pre-test contrast sensitivity are shown using a Cumming estimation plot. Raw contrast threshold data for each participant are plotted on the upper axes; each paired set of observations is connected by a line. On the lower axes, paired Hedges’s *g* is plotted as a bootstrap sampling distribution. Hedge’s *g* value is depicted as dots; 95% confidence intervals are indicated by the ends of the vertical error bars.

For the fellow eyes (**Figure 3** - lower panel), there was a significant interaction between Group and Time, F_3,45_ = 3.303, p = .029, η_p_^2^ = .191. No other omnibus main effects or interactions were significant. During day 1, the active group exhibited a significant improvement in contrast sensitivity from baseline for the post 5 min (*g* = .639 [.127, 1.248], p = 0.033) and post 30 min (*g* = .846 [.199, 1.661], p = 0.018) measurements. No significant differences between baseline and any post-test were found for days 2-5. No significant differences between baseline and any post-test were found within the sham group for any day.

## Uncrowded Visual Acuity

For the amblyopic eyes (**Figure 5** - upper panel), there was a significant interaction between Group and Time, F_3,45_ = 3.325; p = .029, η_p_^2^ = .192). No other omnibus main effects or interactions were significant. During day 1, the active group exhibited a significant improvement in uncrowded visual acuity from baseline for all post-test measurements (during: *g* =.224 [.084, .575], p=.010; post 5: *g =* .281 [.009, .640], p= .05; post 30: *g* = .307 [.118, .795], p = .003). During days 2 and 3, the active group exhibited a significant difference between baseline and only the post 5 min measurement (day 2: *g* = .231 [.091, .383], p = .015, day 3: *g* = .126 [.003, .304], p = .038). No significant differences between baseline and any post-test were found during days 4 and 5. No significant difference between baseline and any post-test was found within the sham group for any day. By chance, there was a substantial difference in baseline uncrowded Landolt-C visual acuity between the active and sham group (compare the dashed lines in **Figure 5** - upper panel).

**Figure 5:**
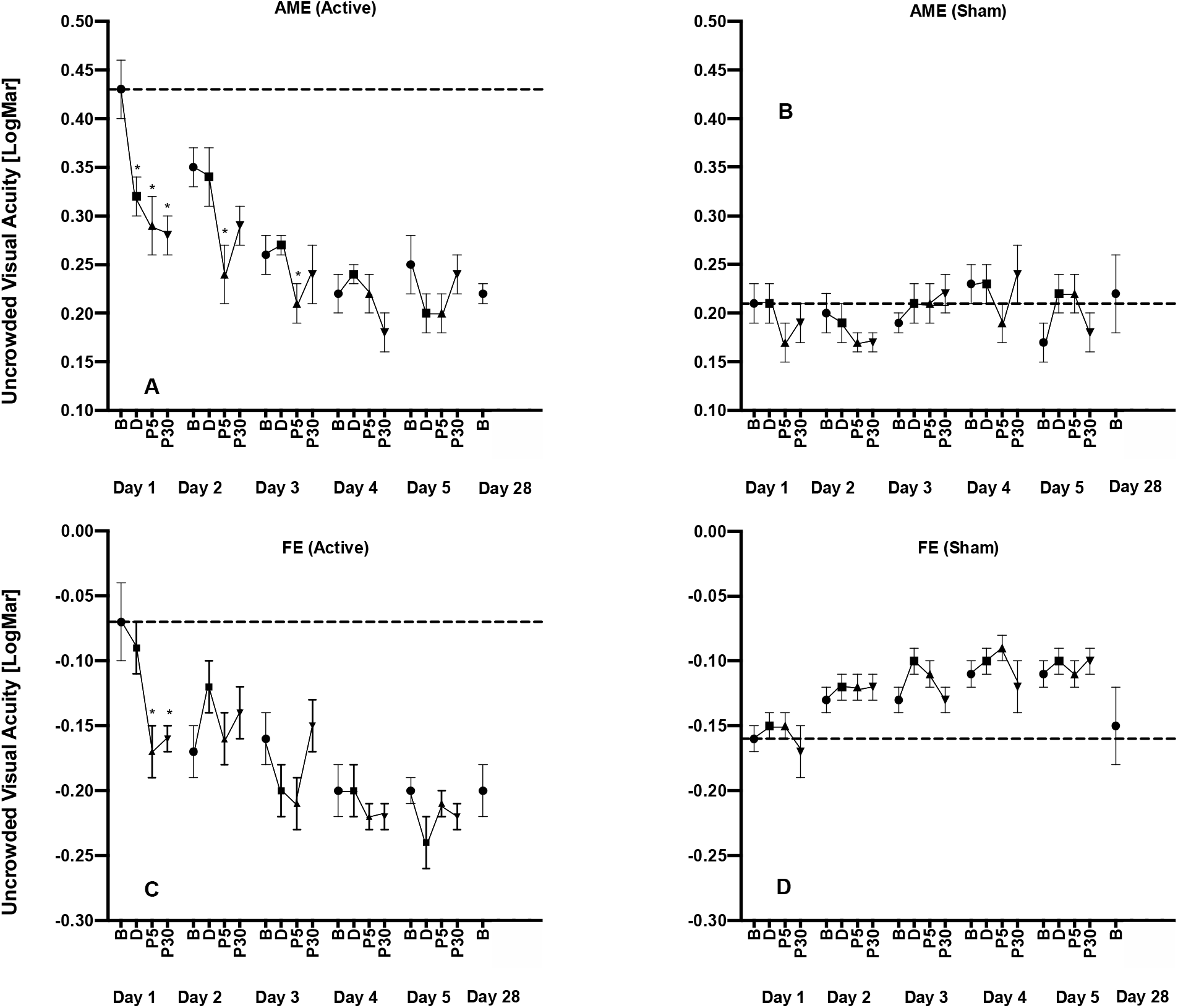
The effects of tRNS on uncrowded visual acuity during each daily session and at the day 28 follow-up visit. Data are shown as in Figure 3. Lower (smaller/more negative) y-axis values indicate better uncrowded visual acuity.

For the fellow eyes (**Figure 5** - lower panel), there was a significant interaction between Group and Time, F_3,45_ = 3.504; p = .023, η_p_^2^ = .200. No other omnibus main effects or interactions were significant. During day 1, the active group exhibited a significant improvement in contrast sensitivity from baseline for the post 5 (*g* = .817 [.164, 1.75], p = 0.035) and post 30 (*g* = .774 [.199, 1.54], p = 0.02) min measurements. No significant differences between baseline and any post-test were found for days 2-5. No significant differences between baseline and any post-test were found within the sham group for any day.

### Crowded Visual Acuity

For the amblyopic eyes (**Figure 6** - upper panel), there were no significant main effects or interactions (all p > 0.05). No significant changes from baseline were observed for any day for any group. As for uncrowded visual acuity, there was a substantial difference in baseline performance between the two groups that occurred by chance during randomization.

**Figure 6:**
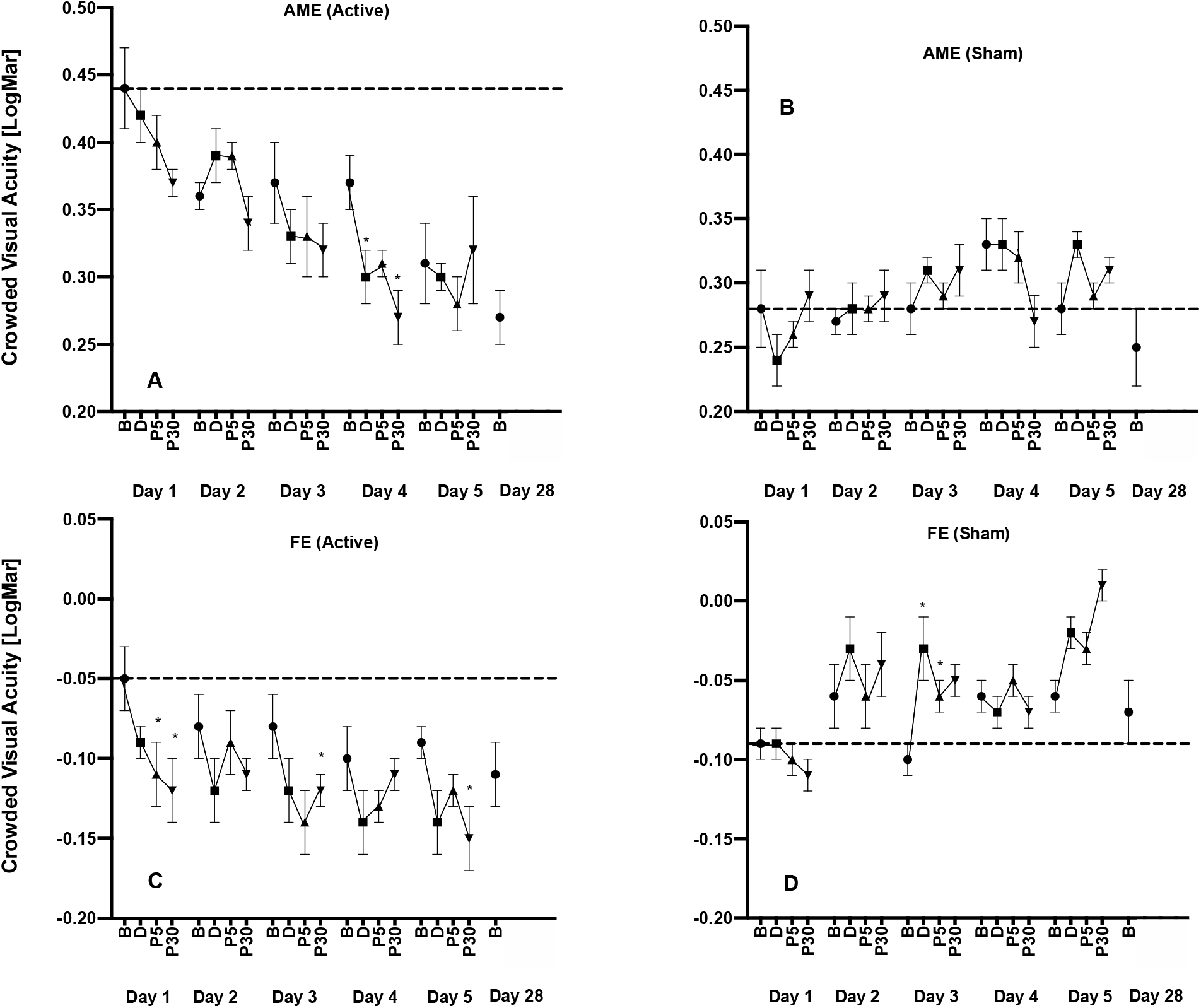
The effects of tRNS on crowded visual acuity during each daily session and at the day 28 follow-up visit. Data are shown as in Figure 3. Lower (smaller/more negative) y-axis values indicate better crowded visual acuity.

For the fellow eyes (**Figure 6** - lower panel), there was a significant interaction between Group and Time, F_3,45_ = 5.733; *p* = .002, *η*_*p*_^*2*^ = .291. No other omnibus main effects or interactions were significant. During day 1, the active group exhibited a significant improvement in crowded acuity from baseline for the post 5 (*g* = .404 [.083, .9], p = 0.05) and post 30 (*g* = .457 [.09, .913], p = 0.039) min measurements. During days 3 and 5, the active group exhibited a significant improvement in crowded acuity from baseline to post 5 min (*g* = .389 [.065, 1.06], *p = .007)* and post 30 min (*g* = .721 [.047, 1.4], *p = .044*) respectively. No significant differences between baseline and any post-test were found for days 2 and 4. No significant differences between baseline and any post-test were found for the sham group.

### Cumulative and long-term effects of tRNS

For the amblyopic eyes, there was a significant interaction between Group and Baseline for uncrowded visual acuity (F_5,65_ = *3.372; p* = *.009*, *η*_*p*_^*2*^ = .206). Pairwise comparisons for the active group revealed a significant difference between the day 1 baseline and the day 3 (*g* = .372 [.163, .771], p = 0.011), day 4 (*g* = .461 [.243, .93], p < 0.003), day 5 (*g* = .369 [.065, .809], p = 0.034), and day 28 (*g* = .454 [.219, 1.03], p = 0.003) baselines. However, no pairwise comparisons were significant for the sham group. There were no significant interactions for amblyopic eye contrast sensitivity or crowded visual acuity measurements or any of the fellow eye measurements.

## Discussion

Our results partially supported our experimental hypothesis that five daily sessions of visual cortex tRNS would improve amblyopic eye contrast sensitivity as well as crowded and uncrowded visual acuity in adult patients. We observed tRNS-induced improvements in contrast sensitivity and uncrowded visual acuity relative to the sham group for both amblyopic and fellow eyes. Crowded visual acuity improved for the fellow but not the amblyopic eyes. Across all outcome measures, pairwise comparisons revealed that acute tRNS effects were statistically significant on day 1 but became non-significant for later sessions. Only amblyopic eye uncrowded visual acuity exhibited a lasting effect of tRNS at follow-up. Our discussion will focus primarily on the results for contrast sensitivity because initial baseline performance was matched between the groups. There were pronounced between-group baseline differences for amblyopic eye uncrowded and crowded visual acuity that occurred by chance during the randomization procedure (randomization occurred before baseline measures were conducted). The difference in baseline performance for the acuity outcome measures make it difficult to properly segregate tRNS effects from task learning effects. One reason for these baseline differences might be a small difference in the proportion of patients with anisometropic amblyopia in active (78%) and sham (60%) groups. However, a much larger scale study will be required to determine whether amblyopia subtype influences the response to visual cortex tRNS.

### tRNS-induced improvements in contrast sensitivity

Our observation that visual cortex tRNS improved amblyopic eye contrast sensitivity is consistent with a growing literature reporting improved contrast sensitivity, visual acuity, stereopsis, and an enhanced cortical response to amblyopic eye inputs following non-invasive visual cortex stimulation in adults with amblyopia [26,37–40,66,67]. Two potential mechanisms have been proposed for tRNS effects: stochastic resonance and changes in the resting membrane potential [51,68]. Stochastic resonance refers to an improvement in signal to noise ratio when a certain amount of noise (in this case neural noise induced by tRNS) is added to non-linear systems [50]. A number of psychophysical studies have provided compelling evidence that stochastic resonance occurs during visual cortex tRNS [24,69–71]. It is possible that the during-stimulation improvements we observed on day 1 for amblyopic eye contrast sensitivity were due to stochastic resonance. However, tRNS aftereffects (i.e. effects that outlast the duration of stimulation) cannot easily be explained by stochastic resonance. Rather, aftereffects may be related to repeated opening and closing of sodium channels during stimulation that causes neural membrane depotentiation and increased cortical excitability. In agreement with this theory, blocking sodium channels appears to reduce the aftereffect of tRNS on motor cortex excitability [72].

A previous study [36] reporting improved amblyopic eye contrast sensitivity following both excitatory and inhibitory visual cortex rTMS proposed a mechanism linked to cortical homeostasis. According to this hypothesis, excitatory stimulation has a more pronounced effect on weakly activated/suppressed neural populations whereas inhibitory stimulation has a greater effect on strongly activated populations. Therefore, both excitatory and inhibitory stimulation are capable of restoring a level of homeostasis to the amblyopic visual cortex by reducing the difference in activation between amblyopic eye dominated neurons (weak activation/suppression) and fellow eye dominated neurons (strong activation) [39]. This, in turn, reduces suppression and/or the relative attenuation of amblyopic-eye-driven neural activity. It is plausible that the excitatory tRNS we employed in this study acts through a homeostatic mechanism.

We also observed improved fellow eye contrast sensitivity in the tRNS group relative to the sham group. Non-invasive visual cortex stimulation studies have reported varying fellow eye effects. Studies using inhibitory stimulation protocols (1 Hz rTMS and continuous theta burst stimulation; cTBS) have reported reduced fellow eye contrast sensitivity [35, 36] whereas those using excitatory protocols (anodal tDCS and tRNS) [24,40], including the present study, observed improvements. This pattern of results is consistent with the homeostasis hypothesis which predicts relatively impaired fellow eye function following inhibitory stimulation and does not rule out improved fellow eye function following excitatory stimulation. This is because excitatory effects may still occur within neuronal populations dominated by the fellow eye, just to a lesser extent than those dominated by the amblyopic eye.

### Successive and cumulative tRNS effects on contrast sensitivity

A day by day analysis of the contrast sensitivity data revealed that tRNS effects were pronounced for both eyes on day 1. However, the within-session tRNS effects waned across sessions, becoming non-significant by day 2 for both eyes. This reduction in within-session tRNS effects was accompanied by stable session to session baseline performance indicating the absence of a cumulative tRNS effect on contrast sensitivity. The waning of within-session effects is consistent with Clavagnier et al’s (2013) study of repeated cTBS sessions in amblyopia, however cTBS did induce cumulative effects that improved baseline performance across sessions. One possible explanation for the waning of within-session tRNS effects and the absence of a cumulative effect on contrast sensitivity relates to stimulation intensity. The relationship between tRNS intensity and visual function improvement during stimulation is an “inverted U”, whereby stimulation that is weaker or stronger than an optimum level has limited effects [51,52]. Lasting changes in cortical excitability induced by prior sessions of tRNS might shift the optimal stimulation intensity towards lower levels, causing a waning of tRNS effects across sessions if stimulation intensity remains constant. If this is the case, tapering stimulation intensity across sessions would be a possible solution.

Another possible explanation for an effect on day 1 and not subsequent days is a placebo effect. Although this cannot be completely ruled out, the use of a single masked, between subjects design combined with automated collection of outcome measures was intended to minimize this source of bias. In addition, participant reported sensations did not differ significantly between the two groups suggesting the adequate masking was preserved throughout the study.

### tRNS effects of crowded and uncrowded visual acuity

It is not possible to draw strong conclusions relating to the amblyopic eye datasets for crowded and uncrowded visual acuity because there were large between-group differences in baseline performance that occurred by chance. However, baseline group differences were minimal for the fellow eye datasets and the results followed those for contrast sensitivity very closely; significant differences between groups that were characterized by within-session improvements early in the experiment and a gradual waning of tRNS effects. This suggests that transient tRNS effects occur for a range of visual functions and that long lasting effects may occur for uncrowded visual acuity.

#### Study limitations

The primary limitation of this study is the relatively small sample size. However, there is no indication in our data that the lack of long-term effects of visual cortex tRNS on amblyopic eye contrast sensitivity is due to insufficient statistical power. The small sample size did preclude the use of stratification for amblyopia subtype and baseline clinical characteristics within our randomization procedure and this likely contributed to the between group differences in baseline amblyopic eye visual acuity that are present in our data.

## Conclusions

tRNS can induce short-term contrast sensitivity improvements in adult amblyopic eyes, however multiple sessions of tRNS do not lead to enhanced or long-lasting effects. In agreement with previous non-invasive brain stimulation studies, these results demonstrate considerable short-term plasticity within the visual cortex of human adults with amblyopia and suggest that multisession tRNS alone does not lead to long lasting improvements in vision.

## Acknowledgements

This work was supported by CIHR grant 390283, CFI grant 34095 and NSERC grants RPIN-05394 and RGPAS-477166 to BT. The authors have no conflicts of interest to declare.

